# CD56/NCAM mediates cell migration of human NK cells by promoting integrin-mediated adhesion turnover

**DOI:** 10.1101/2023.11.21.567714

**Authors:** Amera L. Martinez, Michael J. Shannon, Tyler Sloan, Emily M. Mace

## Abstract

Natural killer (NK) cells patrol tissue to mediate lysis of virally infected and tumorigenic cells. Human NK cells are typically identified by their expression of neural cell adhesion molecule (NCAM, CD56), yet, despite its ubiquitous expression on NK cells, CD56 remains a poorly understand protein on immune cells. CD56 has been previously demonstrated to play roles in NK cell cytotoxic function and cell migration. Specifically, CD56-deficient NK cells have impaired cell migration on stromal cells and CD56 is localized to the uropod of NK cells migrating on stroma. Here, we show that CD56 is required for NK cell migration on ICAM-1 and is required for the establishment of persistent cell polarity and unidirectional actin flow. The intracellular domain of CD56 (NCAM-140) is required for its function, and the loss of CD56 leads to enlarged actin foci and sequestration of phosphorylated Pyk2, accompanied by increased size and frequency of activated LFA-1 clusters. Together, these data identify a role for CD56 in regulating human NK cell migration through modulation of actin dynamics and integrin turnover.

## Introduction

Expression of CD56 (NCAM) marks human natural killer (NK) cells and uniquely demarcates human NK cell subsets (1, 2). CD56/NCAM is a member of the Ig superfamily that has been best characterized on neurons and plays roles in cell adhesion, axon guidance, and synaptic function (3). CD56/NCAM is commonly expressed in three primary isoforms. While the extracellular domain is conserved between all its isoforms, the NCAM-120 isoform has a GPI anchor and NCAM-140 and −180 isoforms have intracellular domains of varying length. Signaling from CD56 has been poorly characterized in lymphocytes, however in neurons each of the NCAM isoforms can influence downstream signaling through direct and indirect binding. Signaling pathways downstream of NCAM in neurons include a constitutive association of NCAM-140 with p59^fyn^ in axonal growth cones which leads to recruitment of focal adhesion kinase (FAK) upon NCAM cross-linking (4). This pathway is of particular interest given the expression of the FAK homologue Pyk2 in human NK cells and decreased Pyk2 phosphorylation in CD56-knockout (KO) NK cells (5). Expression of CD56 on human NK cell lines promotes cytotoxic function and cytokine secretion, and previous studies have identified decreased Pyk2 phosphorylation resulting from deletion of CD56 on human NK cell lines (5). Cross-linking with anti-CD56 antibodies on NK cells promotes signaling in cytokine activated NK cells, further linking it to functional outcomes mediated by intracellular signaling (6).

In addition to its role in cytotoxicity, CD56/NCAM is implicated in the migration and differentiation of human NK cells (7). Primary human NK cells undergo spontaneous cell migration on stromal cell monolayers used to support NK cell differentiation from hematopoietic precursors (7–9). These stromal cells are often a heterogeneous mix of endothelial and mesenchymal cells that express cell adhesion ligands including VCAM-1 and CD44 and secrete extracellular matrix components, including collagen (8, 10). Deletion of CD56/NCAM on primary NK cells or the NK92 NK cell line impairs cell migration on EL08.1D2 stromal cells (7), however the ligands mediating interactions between NK cells and stromal cells have not been elucidated. In addition to ligands for integrins, EL08.1D2 cells express NCAM, and homotypic (NCAM-NCAM) interactions in trans can induce signaling and cell migration in neuronal cells (11). It has remained unclear whether the previously described interactions between NK cells and stromal cells were mediated by NCAM-NCAM adhesion or by other receptor-ligand interactions. Here, we sought to measure cell migration on ICAM-1, a well-defined integrin ligand of LFA-1, to interrogate the role of CD56/NCAM in NK cell migration independently of NCAM-NCAM interactions.

Lymphocyte cell migration is amoeboid motility regulated by cell adhesion, activation of actomyosin contractility, and actin remodeling, coupled with cell polarity (12). While lymphocytes initially undergo symmetrical spreading on ICAM-1, the continuous actin remodeling that occurs on ICAM-1 coated surfaces, even in the absence of a chemokine gradient or shear flow, leads to symmetry breaking and cell migration in both T cells and NK cells (13–15). Cell polarity in migrating lymphocytes includes the formation of a leading edge and a trailing edge, or uropod. We have previously shown that CD56/NCAM localizes to the uropod of human NK cells migrating on stromal cells (7). Uropod localization is not unusual for adhesion receptors and glycoproteins, and other proteins including CD44 and VLA-4 (integrin *α*4β1) are similarly localized and play a role in signaling for cell migration and polarity (16). The FAK homologue Pyk2 is also localized to the trailing edge of migrating NK cells and translocates to the immune synapse upon target cell contact (17). Pyk2 has multiple sites of tyrosine phosphorylation, including the autophosphorylation site Y402, and phospho-Pyk2 Y881 is predominantly found in the uropod and mediates integrin detachment from substrates to enable T cell migration (18).

Here, we sought to better understand the underlying mechanism for impaired cell migration in CD56-deficient human NK cells. Given the role of LFA-1 in promoting cell polarization and spontaneous NK cell migration (13), we studied spontaneous NK cell migration on immobilized ICAM-1 surfaces by confocal and structured illumination microscopy of NK92 cell lines. Through careful dissection of the effect of CD56 deletion in NK92 cells, including the use of NK cell lines expressing actin reporters, we identify new roles for CD56/NCAM in regulating actin and integrin function in migrating lymphocytes.

## Results

### Localization of CD56/NCAM in human natural killer cells

CD56/NCAM has been previously described to be found in the uropod in primary human NK cells migrating on stromal cells (7), however its spatial localization in relation to other proteins associated with cell migration has not been fully explored. To investigate the cellular localization of CD56/NCAM further, we performed fixed cell confocal microscopy of primary NK cells and the human NK cell line NK92 spontaneously migrating on ICAM-1 coated surfaces. As previously described for cells migrating on stromal cell monolayers (7), CD56/NCAM was found in the uropod marked by enrichment of CD44 in primary NK cells and NK92 cells (Fig 1A, B). While less intense than the localization within the uropod, we also found CD56 localized at the membrane in regions of actin enrichment, including at the leading edge of migrating cells (Fig 1A, B). Line profiles underscored the association between CD56/NCAM and the uropod and leading edge of the cell (Fig. 1C), whereas CD44 was exclusively localized to the cell uropod (Fig. 1D).

**Fig. 1.**
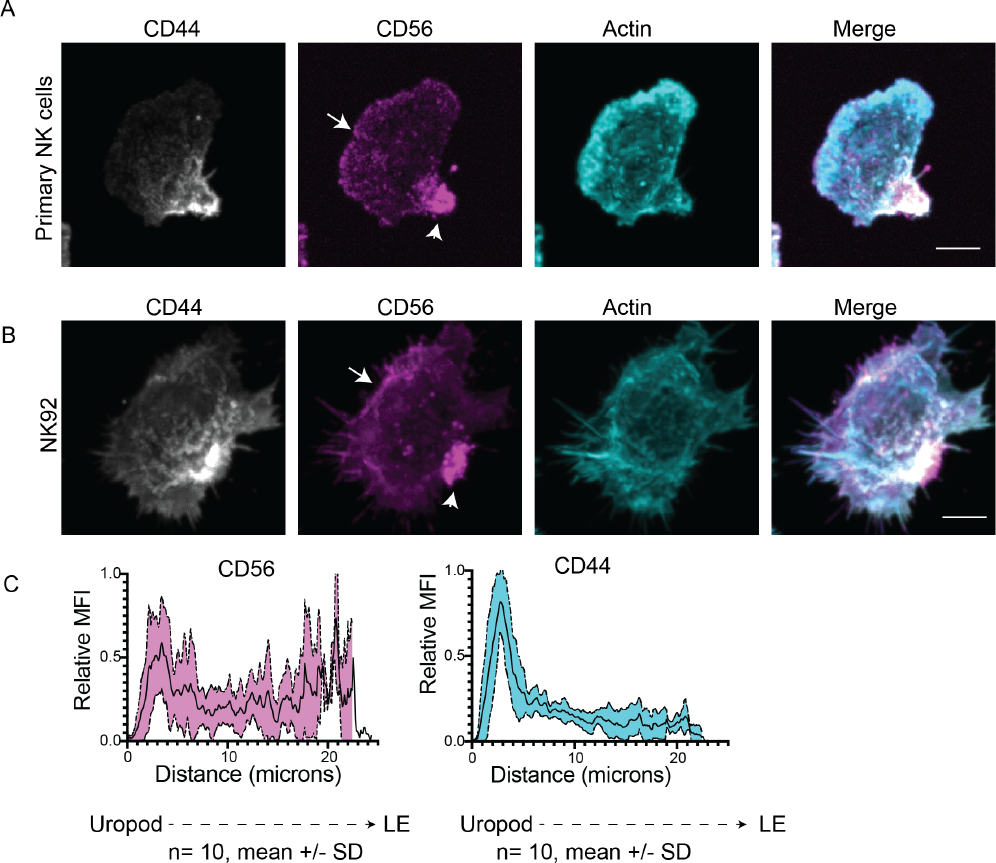
Localization of CD56/NCAM with actin and in the uropod of migrating human NK cells. A) Primary human NK cells undergoing spontaneous migration on immobilized ICAM-1 were fixed, immunostained and imaged by confocal microscopy as indicated. Representative image from 3 biological replicates. B) NK92 cells undergoing spontaneous migration on immobilized ICAM-1 were fixed, immunostained and imaged by confocal microscopy as indicated. C) Representative quantification of the MFI of CD56/NCAM (magenta) or CD44 (cyan) along the cell axis from the cell posterior (uropod) to the leading edge. Mean±SD (SD shown as shaded region) of 10 NK92 cells from one experiment representative of 3 independent experiments. Scale bars 5 µm.

### CD56/NCAM deletion impairs spontaneous cell migration on ICAM-1

Deletion of CD56 in the NK92 cell line impairs NK cell migration on EL08.1D2 stromal cells (7), however the adhesion receptors that mediate spontaneous migration on stroma are not well-defined and could include NCAM itself (5). To evaluate cell migration on isolated integrin ligand and exclude binding to NCAM as a mechanism for CD56-mediated migration, we incubated wild-type and CD56-KO NK92 cells on Fc-ICAM-1 coated surfaces to induce spontaneous crawling migration. Cells were imaged by live cell phase contrast microscopy at 20X magnification using a large field of view. As manual tracking of many cells is time-consuming and can lead to biased measurements, we segmented cells using Cellpose (19) and linked cells between frames using Bayesian tracker (20) (Fig 2A, B). Unlike WT NK92 cells, CD56-KO NK92 cells had impaired migration, yet appeared dynamic and could be seen forming cell protrusions while failing to undergo polarized and directional migration (Fig 2A, B, Movies 1, 2). We additionally noted elongated morphologies, including stretched uropods, of CD56-KO NK92 cells that failed to significantly migrate during the time of imaging (arrowheads, Fig. 2B).

**Fig. 2.**
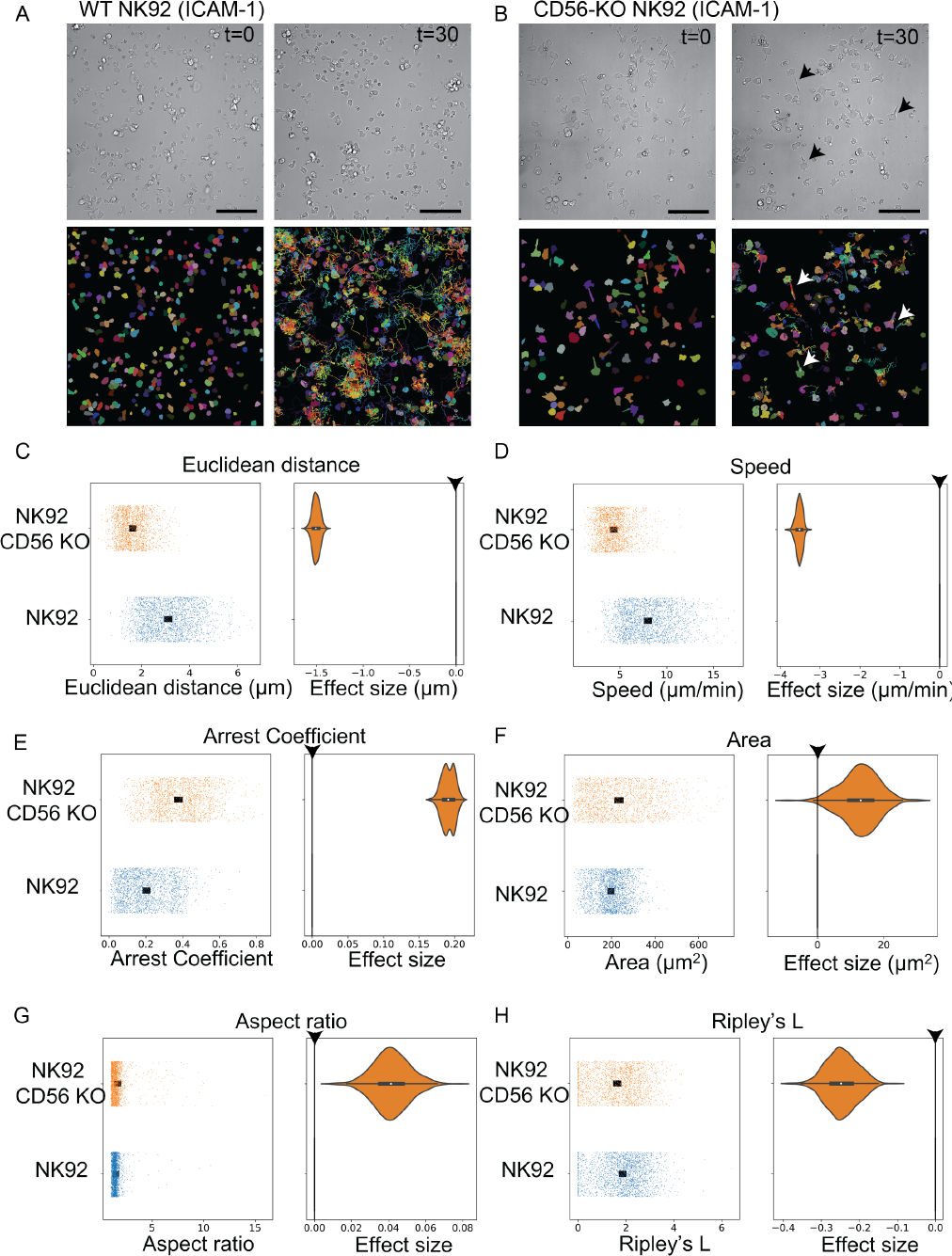
CD56 deletion slows cell migration, decreases cell clustering, and increases cell area. WT or CD56-KO NK92 cells were incubated on ICAM-1 coated glass and imaged by live cell phase contrast microscopy at 10 second intervals for 30 minutes. A-B) Representative micrographs and segmentation by Cellpose that preceded tracking using Bayesian Tracker. C-H) Plots of difference show the distribution of the data (left pane) and effect size (right pane) for metrics related to migration, morphology, and clustering: C) Euclidean distance, D) speed, E) arrest coefficient, F) area, G) aspect ratio, and H) Ripley’s L (clusteredness). n=5 experimental replicates per condition, n= 3946 cells. Cells included in the analysis were filtered to include at least 2 minutes of tracking and a minimal area of 10 µm^2^ to exclude cell debris. Arrowhead indicates zero line for control condition (WT NK92) on Plots of Difference.

Cell segmentation and tracking were followed by analysis using a new custom software pipeline to measure 24 parameters of cell morphology, migration, and cell clustering (21). Consistent with a role for CD56/NCAM in mediating NK92 cell migration, we found decreased migratory capacity of CD56-KO cells when Euclidean distance and speed were considered (Fig 2C, D) and increased arrest coefficient (Fig. 2E). CD56-KO cells had greater cell area and aspect ratio when compared to WT cells (Fig 2F, G), reflecting the cell stretching described above. In addition, CD56-KO cells formed homotypic cell clusters less frequently as measured by Ripley’s L (r) coefficient (Fig 2H).

Our measurements were complemented by related descriptors of cell and migration features, and we additionally found increased cell perimeter, and decreased mean squared displacement (MSD), maximum distance, and cumulative length in CD56-KO cells relative to WT cells (Supp. Fig. 1). These measurements were consistently stable over the time of imaging, thus confirming that our imaging timescales were appropriately capturing cell behaviors (Supp. Fig 2). To confirm that the spontaneous migration we observed was specific to ICAM-1, we included control surfaces coated with Fc only, poly-L-lysine, or anti-CD18 to induce LFA-1 cross-linking. While anti-CD18 induced cell spreading in both WT and CD56-KO cells, quantified by increased cell area, none of these surfaces induced significant spontaneous cell migration (Supp. Fig. 3).

### CD56-KO NK92 cells have decreased unidirectional persistence of actin flow and fail to maintain cell polarity

To investigate the reduced capacity for cell migration by CD56-KO cells further, we imaged WT and CD56-KO NK92 cells expressing LifeAct mScarlet (22) using higher resolution (100X) live cell confocal microscopy. Confocal imaging of actin in live cells confirmed unidirectional polarization and migration of WT NK92 cells, characterized by a dynamic branched actin network at the leading edge, and revealed the multidirectional formation of actin networks in CD56-KO NK92 cells (Fig. 3A, Movies 3, 4). To quantify this observation, we used particle image velocimetry (PIV) for visualization of actin flow (23). To quantify changes in actin flow directionality, we extracted the kappa parameter from a Von Mises fit of 10 second overlapping time windows of flow. This analysis demonstrated that CD56-KO NK92 cells are not able to maintain a directionally persistent flow in the leading edge (Fig 3B, C). Despite their impaired cell migration (Fig. 3D, and Fig. 2), CD56-KO cells had only slightly decreased actin flow speeds (Fig. 3E) and similar immobile actin fractions (Fig. 3F) as WT NK92 cells. Performing the same PIV analysis on actin flow in NK cells incubated on anti-CD18 functionalized surfaces demonstrated extensive actin remodeling in both WT and CD56-KO cells, however actin remodeling occurs at multiple or extended lamellipodia on anti-CD18 and the cells do not polarize or migrate (Supp. Fig 4, Movies 5, 6). CD56-KO cells on ICAM-1 did exhibit a very slightly reduced directional persistence of actin flow, reflecting disordered actin dynamics, but due to the existence of multiple lamellipodia in the WT cells on this surface, the difference was minimal (Supp. Fig 4). Actin remodeling effects were demonstrated to be specific, as treatment with cytochalasin D significantly reduced actin flow speed in both cell types (Supp. Fig 5) and were constant over the 2-minute timelapse imaging period in all cases (Supp. Fig 6). Therefore, while anti-CD18 induces actin remodeling, ICAM-1 uniquely promotes symmetry breaking and spontaneous cell migration. Further, while persistent unidirectional actin flow and maintenance of the leading edge is dependent on CD56/NCAM function, actin remodeling occurs in both WT and CD56-KO lines. To further validate this observation, we imaged WT and CD56-KO lines in collagen matrices by lattice lightsheet microscopy at high temporal and spatial resolution. Live cell imaging of both WT and CD56-KO Life-Act mScarlet NK92 lines revealed dynamic actin processes of cells in collagen, however whereas WT cells were polarized and exhibited flat lamellar projections, CD56-KO NK92 cells were less polarized and had multidimensional projections (Movies 7, 8).

**Fig. 3.**
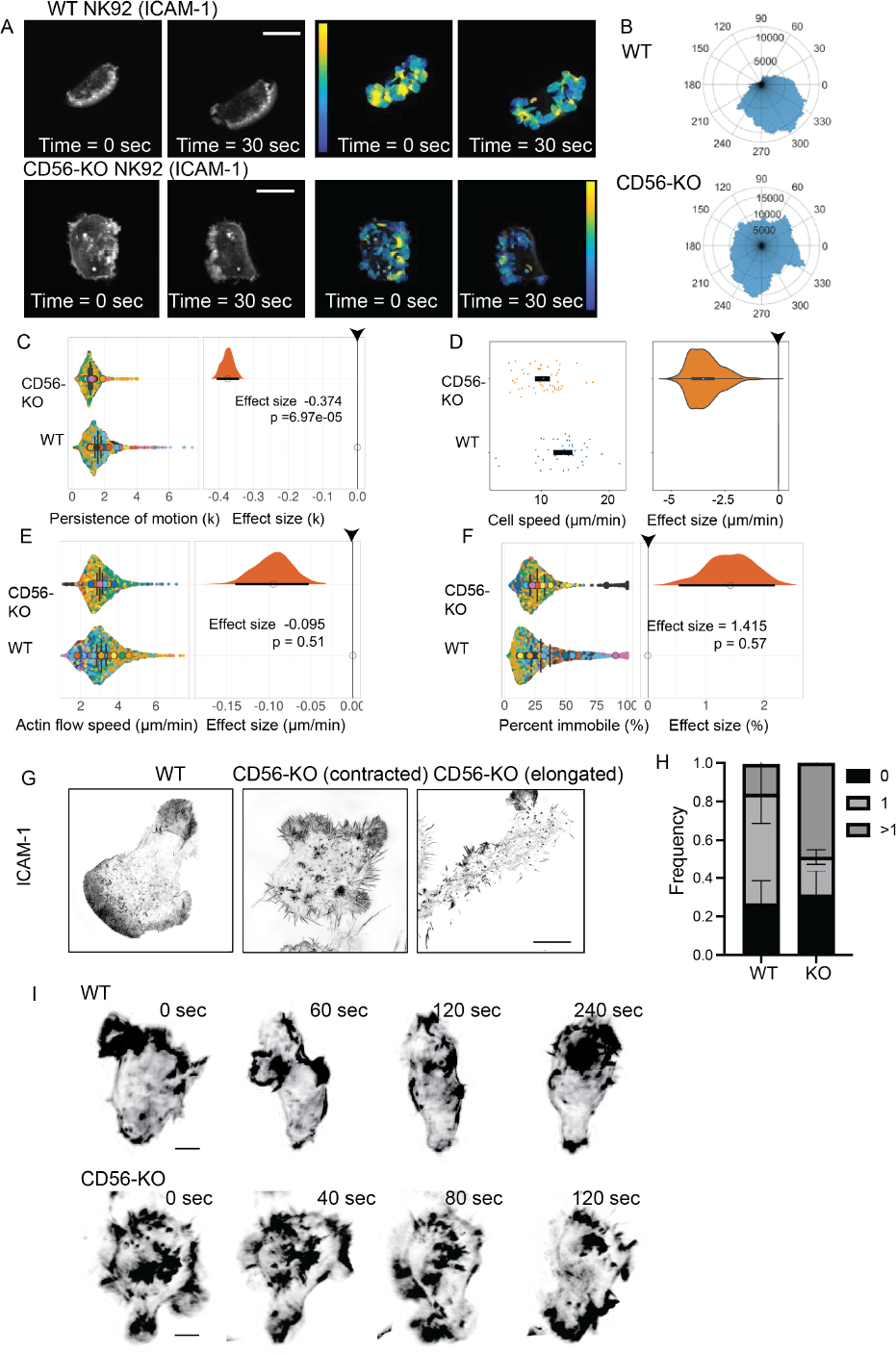
Lamellipodial actin flow polarity is deficient while flow speed is intact in CD56-KO NK92 cells migrating on ICAM-1. WT or CD56-KO NK92 cells expressing LifeAct mScarlet were incubated on ICAM-1 coated glass and imaged by live cell confocal microscopy at 100X magnification for 2 minutes per field of view with 1 second between frames using a 561 nm laser. See also Movies 3 and 4.A) Example frames from a timelapse movie are shown at 0 and 30 seconds (left) with Lucas-Kanade PIV vector field (right); warm colors = high magnitude, cold = low. B) Polar histogram of directionality of PIV vectors. C) Plots of difference showing the concentration of motion of flow (κ from the von Mises fit) across 10 second overlapping time windows illustrating change in flow direction. D) Plots of difference of cell speed of migration. E) Plots of difference of actin flow speed. F) Percent of actin fraction that was immobile. n=30 and 25 experimental replicates per condition respectively, n= 92 cells analyzed. Arrowhead indicates zero line for control condition (WT NK92) on plots of difference (C-F). G) WT or CD56-KO migrating on ICAM-1 were fixed, immunostained for actin (phalloidin) and imaged by SIM. Scale bar 5 µm. H) WT or CD56-KO NK92 cells migrating on ICAM-1 were scored for the presence of 0, 1 or >1 lamellipodia. Mean±SD, n = 150 (WT) and 262 (CD56-KO) from 3 independent experiments. I) Lattice lightsheet imaging of WT or CD56-KO NK92 cells expressing LifeAct mScarlet embedded in 1.6 mg/ml collagen and imaged live every 40 sec. Select frames are shown to highlight cell morphologies; see also Movies 7 and 8. Scale bar 5 µm.

Lastly, we performed structured illumination (SIM) microscopy on cells migrating on ICAM-1 to understand how the fine structure of actin fibers are affected by CD56 deficiency. While WT NK92 cells were polarized with a clear leading and trailing edge, CD56-KO NK92 cells often had multipolar phenotypes, and many were either contracted with no clear polarity or elongated as previously observed in live cell imaging (Fig 3G). Quantification of the frequency of cells that formed single or multiple leading-edge structures showed a significantly greater frequency of CD56-KO NK92 cells with >1 leading edge than WT cells, and a slightly greater frequency of CD56-KO NK92 cells with no apparent leading edge (Fig 3H). Together with cell migration data, these data demonstrate that CD56-KO NK92 cells have impaired cell migration and an inability to maintain a persistent leading edge, but that their capacity for dynamic actin remodeling is not significantly impaired.

### The intracellular domain of CD56 is required for actin homeostasis

We noted the presence of dynamic actin foci in the mid-body of cells in both WT and CD56-KO LifeAct mScarlet NK92 cells incubated on ICAM-1 that were more pronounced in CD56-KO NK92 cells (Movies 3,4). Similar structures were observed in WT and CD56-KO NK92 cells incubated on anti-CD18 antibody, which induces symmetrical spreading but not cell migration (Movies 5, 6). Our SIM imaging also showed that CD56-KO cells migrating on ICAM-1 had larger actin foci and longer filopodia than WT NK92 cells (Fig 3G). We decided to focus on the pronounced actin foci in CD56-KO NK cells undergoing symmetrical spreading to look more closely at these structures in the absence of cell polarity and symmetry breaking (Fig. 4A). 3D reconstruction of these images showed that actin foci in CD56-KO cells were taller in the Z dimension as well as XY (Fig 4B) and further highlighted the presence of filopodia seen in XY imaging at the plane of the glass. Given the distinctive actin structures seen in CD56-KO NK92 cells, we hypothesized that CD56/NCAM may be mediating actin remodeling via its intracellular domain. CD56/NCAM is expressed as 3 common isoforms, and the NCAM-140 isoform is predominantly expressed in human NK cells (1, 2, 5, 24). We reconstituted CD56-KO NK92 cells with full-length NCAM-140 (5) or NCAM-140 lacking its intracellular domain (ΔICD). Flow cytometry confirmed the surface expression of CD56/NCAM in all reconstituted lines (Fig 4C). Probing by Western blot confirmed that the molecular weight of the full-length construct corresponded to WT CD56/NCAM, whereas the truncated isoform had decreased molecular weight predicted by the loss of the intracellular domain (Fig. 4D). To determine whether the intracellular signaling domain of CD56/NCAM-140 is required to mediate actin homeostasis, we incubated CD56-KO cells that had been reconstituted with full-length or truncated (ΔICD) CD56/NCAM on anti-CD18 coated surfaces and measured the accumulation of actin foci using SIM. As shown in Fig 4A, CD56-KO cells had significantly larger actin foci than WT NK92 (Fig 4E). As predicted, cells expressing full-length CD56 had actin foci that were not significantly different in size from WT NK92 cells. Cells expressing the truncated (ΔICD) form of CD56 had actin foci that were enlarged, and actin foci areas in these cells were not significantly different from the CD56-KO condition (Fig 4E). Similarly, actin filopodia length was restored in CD56-KO NK92 cells expressing the full-length 140 kDa isoform, but not ΔICD, of CD56 (Fig 4E). Quantification confirmed the significantly increased size of actin foci and increased length of filopodia, and the rescue of these effects by full-length, but not truncated, CD56/NCAM-140 (Fig 4F). Together these data demonstrate that the role that CD56/NCAM-140 plays in regulating actin phenotypes is dependent on the intracellular domain of NCAM-140.

**Fig. 4.**
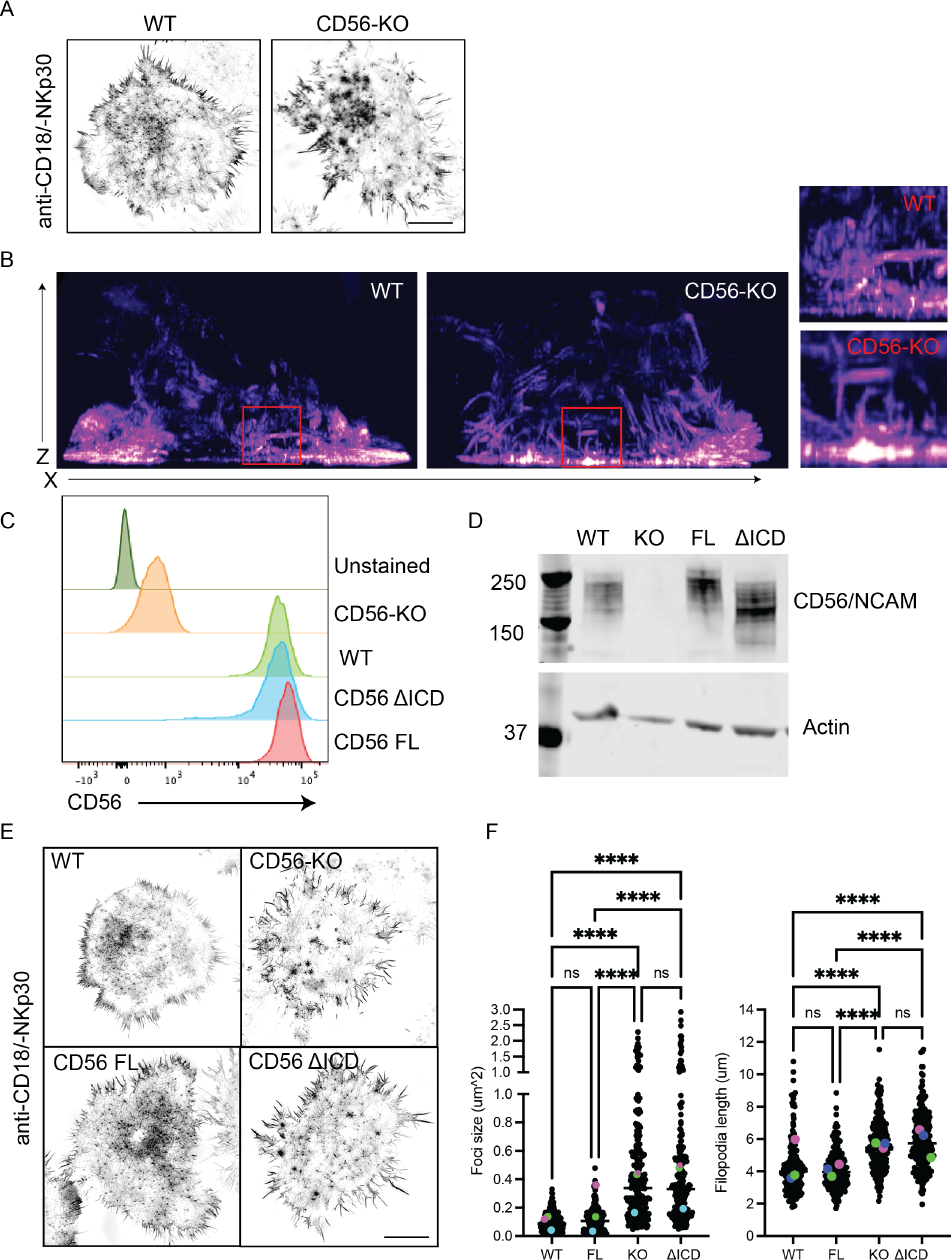
The intracellular domain of CD56 regulates actin and filopodia homeostasis. A) WT or CD56-KO NK92 cells were incubated on anti-CD18/-NKp30 then fixed and immunostained for actin (phalloidin) and imaged by 3D-SIM. Scale bar 5 µm. B) 3D reconstruction and XZ view of cells imaged as in (A). Inset highlights depths of actin accumulation. C) Flow cytometry of cell surface CD56 expression. D) Western blot of CD56/NCAM with actin as a loading control. E) WT or CD56-KO NK92 or CD56-KO reconstituted with full-length or delta ICD CD56 were incubated on anti-CD18/-NKp30 then fixed, immunostained for actin (phalloidin) and imaged by SIM. F) Foci size and filopodia length were quantified for cell lines shown in (E). Foci n=183 (WT), 225 (FL), 229 (KO), 215 (delta ICD). Filopodia n= 192 (WT), 221 (FL), 204 (KO), 206 (delta ICD). Mean from each of 3 independent experiments are highlighted. Scale bar 5 µm.

### Impaired integrin adhesion turnover in CD56-KO cells is associated with changes in Pyk2 Y402 and Pyk2 Y881 localization

Previous studies have shown that CD56 deletion leads to decreased phosphorylation of the protein tyrosine kinase Pyk2 (5) and that Pyk2 can play a role in uropod detachment and lymphocyte migration, particularly through phosphorylation on the Y881 motif (18). As previously reported in CTLs, we found that Pyk2 phospho-Y881 was localized to the uropod of NK92 cells migrating on ICAM-1 coated surfaces (Fig 5A). In contrast, Pyk2 phospho-Y402 was more diffusely localized and there was a smaller pool of Pyk2 phospho-Y402 also found in the uropod (Fig 5A). CD56-KO NK92 cells formed multipolar patterns on ICAM-1 as previously shown, including large actin foci. These foci were highly enriched for Pyk2 phospho-Y402 (Fig. 5A). Line profiles across pPyk2 foci further demonstrated enrichment of actin in Pyk2 phospho-Y402, but not phospho-Y881, foci in CD56-KO NK92 cells (Fig 5B).

**Fig. 5.**
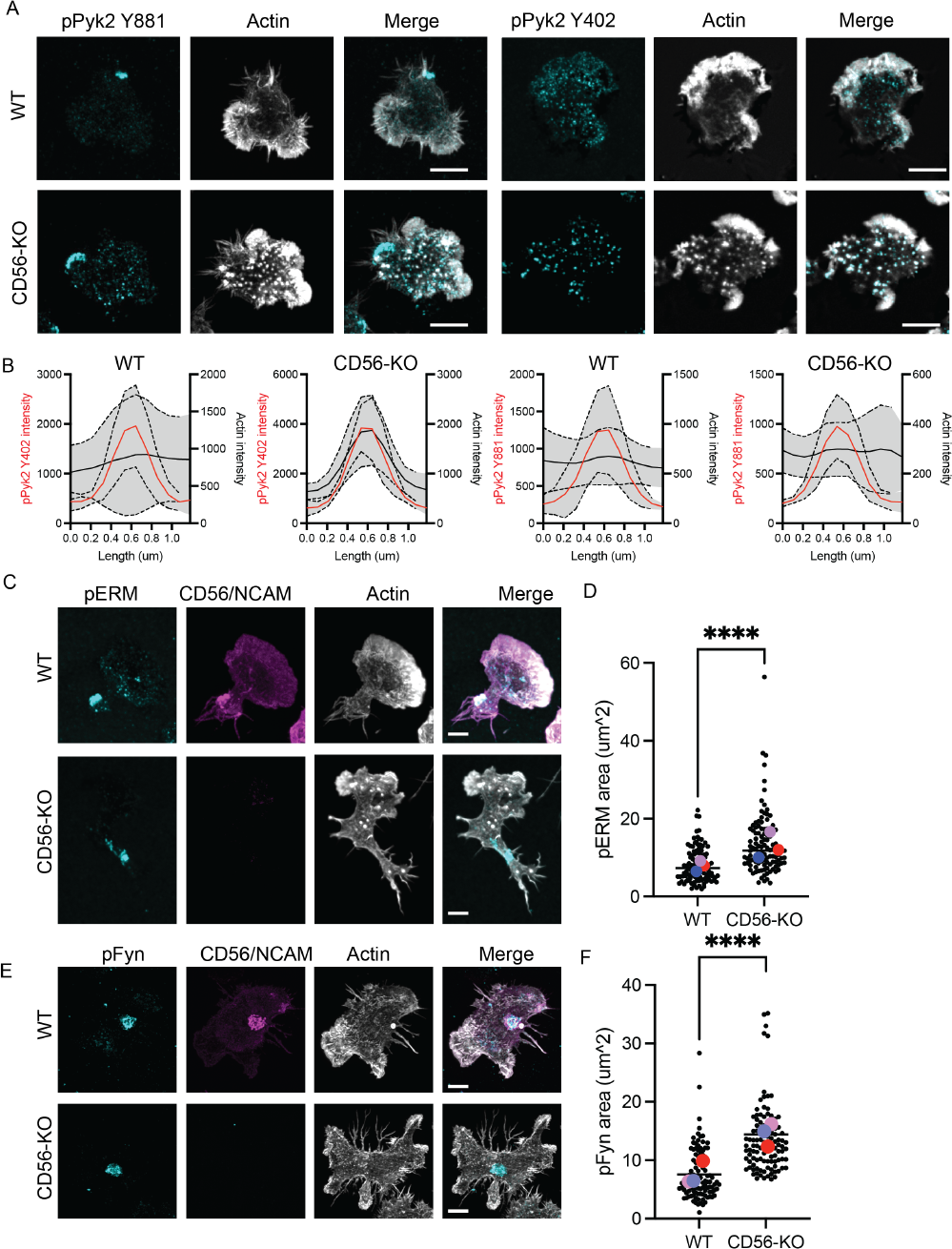
pPyk2 localization is dysregulated in CD56-KO NK92 cells. A) WT and CD56-KO NK92 cells were incubated on ICAM-1 coated surfaces for 45 minutes then fixed and immunostained for actin (phalloidin) and Pyk2 phospho-Y881 or phospho-Y402 as indicated. Representative images from the plane of the glass are shown. B) Line profiles drawn across actin foci in WT or CD56-KO NK92 cells were used to generate actin intensity (black line) are shown with corresponding Pyk2 phospho-Y402 or Pyk2 phospho-Y881 intensity (red line). n=10 foci per condition. Solid line, mean; dashed line standard deviation. C) Cells were treated as in (A) and immunostained for phospho-ERM. Representative maximum projections are shown. D) Quantification of pERM area from maximum projection images shown as Superplots with individual experiments highlighted. n= 107 (WT), 108 (CD56-KO) from 3 independent experiments, large symbols indicate the mean from individual experiments. D) Cells were treated as in (A) and immunostained for phospho-Fyn. Representative maximum projections are shown. D) Quantification of phospho-Fyn area from maximum projection images shown as Superplots with average from independent experiments highlighted. n= 104 (WT), 108 (CD56-KO) from 3 independent experiments, large symbols indicate the mean from individual experiments. Scale bar 5 µm.

The presence of multiple lamellipodial structures in CD56-KO cells suggested that establishment of cell polarity was impaired in CD56-KO cells attempting to migrate on ICAM-1 coated surfaces. Alternatively, the presence of actin foci enriched for phospho-Pyk2 suggested that dysregulation of integrin-mediated adhesion turnover may be the driver of this phenotype. To distinguish between these two mechanisms, we detected phospho-ERM (pERM) proteins, as these are also associated with establishing cell polarity in migrating lymphocytes. In both WT and CD56-KO NK92 cells we found ERM phospho-Thr558 in a localized patch in cells on ICAM-1 (Fig 5C). In WT cells, this patch of pERM was localized in the cell uropod. The presence of multilamellar structures made it more difficult to identify a clear uropod in CD56-KO NK92 cells, however the presence of clustered pERM suggests that initial signaling to generate cell polarity on ICAM-1 is preserved. We did note, however, that the area of pERM detected in CD56-KO NK92 cells was reproducibly greater than in WT NK92 cells, suggesting that CD56 may be playing a role in constraining ERM proteins (Fig 5D).

Finally, in neurons CD56/NCAM is associated with the Src kinase Fyn, which is also expressed in lymphocytes and mediates NK cell effector functions and T cell chemotaxis (25, 26). To determine whether loss of CD56/NCAM affected Fyn localization or phosphorylation, we immunostained for phospho-Fyn in WT and CD56-KO NK92 cells migrating on ICAM-1 (Fig 5E). We found that phospho-Fyn was similarly localized to the uropod of migrating cells, and loss of CD56/NCAM similarly led to increased area of phospho-Fyn in CD56-KO cells (Fig 5F). These observations suggest that polarity induced by ERM protein and Fyn phosphorylation is not disrupted upon loss of CD56/NCAM but that CD56 may be playing a role in constraining polarity proteins following cell polarization.

### Loss of CD56/NCAM leads to an increase in activated LFA-1 on the cell surface

Given the apparent differences in integrin-mediated adhesion and intact cell polarity signaling in CD56-KO cells, we investigated the structure of LFA-1 focal complexes in NK92 cells migrating on ICAM-1. Detection of open conformation LFA-1 by confocal microscopy using the monoclonal antibody m24 identified a greater number of high-affinity LFA-1 foci with a greater area and intensity in CD56-KO NK92 cells on ICAM-1 when compared to WT cells (Fig 6A, B). Reconstruction of 3D images from Z-stacks supported this observation and identified enrichment of m24 staining in the glass-proximal slices in CD56-KO NK92 cells relative to WT (Fig. 6B, right). Flow cytometry confirmed that total cell surface expression of β2 integrin (CD18) was not affected by loss of CD56/NCAM (Fig. 6C).

**Fig. 6.**
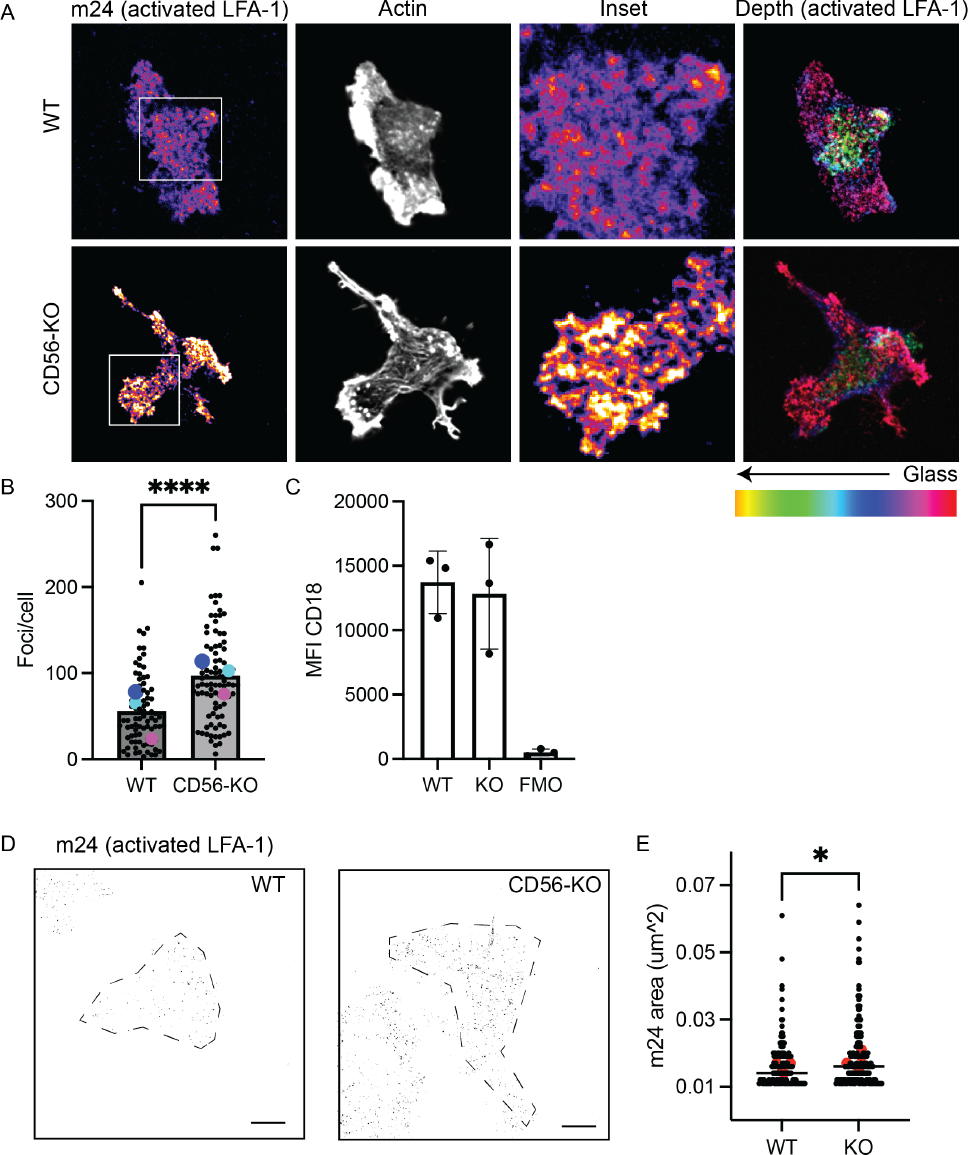
Increased LFA-1 activation in CD56-KO NK92 cells on ICAM-1. A) WT and CD56-KO NK92 cells were incubated on ICAM-1 coated surfaces for 45 minutes then fixed and immunostained for actin (phalloidin) and open conformation LFA-1 (clone m24) as indicated and imaged by confocal microscopy. Depth color shading of Z stacks taken throughout the image is shown to the right, all other images are a single plane of the cell proximal to the glass. Scale bar 5 µm. B) Quantification of the frequency of m24 foci/ per cell shown as Superplots with the average from each experiment highlighted. n= 83 (WT), 90 (CD56-KO) from 3 independent replicates. C) Mean fluorescence intensity of surface expression of integrin β2 (CD18) on WT and CD56-KO NK92 cells measured by flow cytometry; FMO, fluorescence minus one control. n=3 independent technical replicates. D) Visualization of m24 prepared as in (A) and imaged by 3D-SIM. Images shown are maximum project of 3 slices at the plane of the glass. Scale bar 5 µm; cell footprint outlined by dashed line. E) Quantification of m24 foci area from SIM microscopy data. Representative data from 3 independent experiments.

As detection of active LFA-1 foci by confocal microscopy was constrained by the diffraction limit, we additionally imaged active LFA-1 on NK92 cells on ICAM-1 using SIM (Fig 6D). This approach allowed us to measure the area of high affinity LFA-1, which identified a larger size of foci in CD56-KO NK92 cells relative to WT cells (Fig 6E). This observation, together with larger phospho-Pyk2 enriched actin foci in CD56-KO cells, suggests that integrin-mediated adhesions are not being appropriately disassembled in CD56-KO NK92 cells.

Given the apparent differences in integrin-mediated adhesion and intact cell polarity signaling in CD56-KO cells, we investigated the structure of LFA-1 focal complexes in NK92 cells migrating on ICAM-1. Detection of open conformation LFA-1 by confocal microscopy using the monoclonal antibody m24 identified a greater number of high-affinity LFA-1 foci with a greater area and intensity in CD56-KO NK92 cells on ICAM-1 when compared to WT cells (Fig 6A, B). Reconstruction of 3D images from Z-stacks supported this observation and identified enrichment of m24 staining in the glass-proximal slices in CD56-KO NK92 cells relative to WT (Fig. 6A, right). Flow cytometry confirmed that total cell surface expression of β2 integrin (CD18) was not affected by loss of CD56/NCAM (Fig. 6C). As detection of active LFA-1 foci by confocal microscopy was constrained by the diffraction limit, we additionally imaged active LFA-1 on NK92 cells on ICAM-1 using SIM (Fig 6D). This approach allowed us to measure the area of high affinity LFA-1, which identified a larger size of foci in CD56-KO NK92 cells relative to WT cells (Fig 6E). This observation, together with larger phospho-Pyk2 enriched actin foci in CD56-KO cells, suggests that integrin-mediated adhesions are not being appropriately disassembled in CD56-KO NK92 cells.

### Discussion

Previous studies of the role of CD56/NCAM in human NK cell function indicated a role for CD56/NCAM in cell migration on stromal cells. Specifically, CD56/NCAM was found localized to the cell uropod of primary NK cells migrating or arrested on stroma (7). Deletion of CD56/NCAM in the NK92 cell line also impaired cell migration on stroma, suggesting that CD56/NCAM mediates NK-stromal cell interactions to promote motility of NK cells. Our current study advances previous studies of the role of CD56 in NK cell migration by demonstrating that CD56-deficient cells have impaired cell migration on the integrin ligand ICAM-1, and that loss of CD56/NCAM leads to impaired sustained cell polarity and increased detection of activate LFA-1 at the plane of activation on functionalized glass. This phenotype is accompanied by a striking actin cytoskeleton profile including enlarged actin foci and elongated filopodia present following ICAM-1 ligation by LFA-1 or integrin activation by antibody cross-linking. CD56/NCAM-deficient NK92 cells migrating on ICAM-1 had intact actin flow speed and amount of immobile actin but actin remodeling was not maintained with persistent directionality, resulting in multidirectional, short lived lamellipodia. While we have previously shown that adhesion to target cells is unaffected in CD56-KO NK92 cells (5), directionality of actin flow in CD56-KO cells was highly irregular while flow speed was the same, indicating the significant role of actin dynamics in impaired cell migration in the absence of CD56. Further, establishment of cell polarity as indicated by ERM phosphorylation and capping was also largely intact, suggesting that the root cause of impairment in forward cell migration may be further downstream. Interestingly, the actin phenotype that we observed, namely large foci in CD56-deficient cells, was also present in cells which had undergone symmetrical spreading, indicating that integrin turnover is occurring in the absence of forward cell movement.

While phosphorylation of Pyk2 is globally decreased in CD56-KO NK cells (5), we now demonstrate that this global decrease in Pyk2 phosphorylation is associated with increased segregation of Pyk2 phospho-Y402 in enlarged actin complexes found in CD56/NCAM-KO cells. We further demonstrate that, as in T cells (18), Pyk2 phospho-Y881 in migrating human NK cells preferentially localizes to the cell uropod, where it likely plays a role in mediating detachment from ICAM-1 coated surfaces. The mechanism by which NCAM regulates the interplay between integrin, actin, and cell migration is still unclear. The phenotype that we observe, namely of multiple leading edges formed as cells try to migrate but are unable to detach their uropod, is highly reminiscent of T cells treated with Pyk2 inhibitor or dominant negative C-terminal FAT domain of Pyk2 (18). Together with previous reports showing Pyk2 phosphorylation in response to CD56 cross-linking in human NK cells (6), this data supports a role for CD56/NCAM in mediating signaling through Pyk2, including uropod detachment. Pyk2 deficient macrophages have impaired chemokine-induced migration that includes impaired trailing edge detachment, further implicating Pyk2 function in cell detachment during migration and generating a cell phenotype resembling that seen in CD56-deficient NK cells (27). In neurons, CD56/NCAM binds directly to the Src family kinase member Fyn through the NCAM-140 intracellular domain and mediates FAK signaling in lipid rafts (4, 28). Here we show that the intracellular domain of CD56/NCAM is required for its function, suggesting that steric hindrance by polysialated CD56/NCAM on the cell surface is not primarily the mechanism by which CD56/NCAM regulates integrin function. CD56/NCAM may act similarly in lymphocytes to regulate interactions between Fyn, Pyk2, and integrins which are required for actin homeostasis.

While actin flow speed in CD56-KO NK92 cells is minimally affected, CD56-KO NK92 cells form multipolar phenotypes in response to integrin ligation and demonstrate impaired uropod retraction and forward movement. This phenotype suggests that the actin phenotype that we observe, namely enlarged actin clusters and long filopodia, results from reduced Pyk2 phosphorylation and Pyk2 mislocalization. Actin regulation requires the intracellular domain of CD56/NCAM, suggesting that it is not simply steric modulation via polysialated CD56/NCAM. However, whether CD56/NCAM can play a similar role in the arrangement of other cell surface receptors remains to be seen and may be related to previously reported defects in cytotoxic function and cytokine secretion by CD56-deficient human NK cells. Recent studies of CD44 have identified its role in modulating the macrophage glycocalyx by acting as a picket protein (29), and it is conceivable that CD56/NCAM could serve a similar function. Similar studies have identified integrins as subject to glycocalyx regulation (30), however single molecule studies of CD56/NCAM, Pyk2 and integrins are required to fully elucidate the relationship between them.

The function of CD56/NCAM has been difficult to fully elucidate, likely in part because it is not acting in conventional receptor-ligand interactions. Regulation of integrin detachment enables cell migration on 2-dimensional surfaces, and impaired integrin inactivation or integrin endocytosis following ligand binding prevents cell migration (31–33). While our studies were primarily performed on ICAM-1 coated surfaces, there is no indication that CD56/NCAM is directly binding to ICAM-1, as large actin foci and elongated filopodia were also present in response to anti-CD18 cross-linking. The use of isolated ligands in the current study also avoids the potential binding of CD56/NCAM to other NCAM molecules present on stromal cells previously used to measure NK cell migration (7), convincingly ruling out NCAM-NCAM interactions as the mediators of CD56-dependent cell migration on stromal cells. While not previously shown in NK cells, cell adhesion molecules can mediate integrin detachment and turnover. L1CAM, which is structurally similar to NCAM, promotes endocytosis of integrin β1 to enable forward cell migration (34). Whether CD56/NCAM promotes appropriate initial integrin binding and receptor engagement or functions solely in detachment via deactivation or endocytosis, remains to be seen.

Many interesting questions remain about the biological function of CD56/NCAM, including the way in which human NK cell subsets with different densities of CD56/NCAM undergo cell migration and the single-molecule dynamics of CD56 during migration. In addition to differences in CD56 density, human NK cell subsets have extensive differences in actin and integrin machinery (35). While acute loss of CD56/NCAM in NK cell lines with high CD56 expression, such as NK92 cells, leads to impaired cell migration, closer study of its role in other NK cell subsets will be necessary to fully understand how it interacts with other signaling pathways that regulate cell migration. Similarly, the study of other immune cells that express CD56/NCAM, including NKT cells, myeloid precursors in mice, and transformed cells such as myeloma, remains an interesting area of further study. Finally, it must be noted that the current studies were performed on 2-dimensional surfaces in the absence of shear flow, and it will be of interest to further understand how CD56/NCAM-deficient cells interact with 3-dimensional and complex microenvironments. Nevertheless, our current study sheds light on the cell biological function of a ubiquitously expressed but poorly understood receptor on human immune cells and its function in cell migration.

## Methods

### Cell lines and cell culture

CD56-KO cells were generated and described previously (7). WT, CD56-KO and LifeAct-mScarlet NK92 cells were maintained in culture in MEM alpha medium without ribonucleosides and deoxyribonucleosides but with 2 mM L-glutamine and 1.5 g/L sodium bicarbonate (Gibco; 12-561-056), supplemented with 12.5% fetal bovine serum (FBS), 12.5% horse serum, 100 U/ml IL-2 (ProLeukin; 65483-116-07), 200 µM myoinositol (Thermo J62886.22), 20 µM folic acid (Sigma F8758-25G), 100 µM 2-mercaptoethanol (Gibco; 21985-023) and 1% Pen-Strep antibiotics. pLifeAct mScarlet N1 was a gift from Dorus Gadella (Addgene plasmid 85054; RRID:Addgene 85054). WT and CD56-KO NK92 cells were nucleofected with pLife-Act mScarlet using an Amaxa nucleofector (Kit R) and cells were selected with G418 before validating expression by microscopy and flow cytometry.

### Flow cytometry

Flow cytometry was performed on cell lines as independent technical replicates with different passages of cells on different days. NK92 cells were harvested, washed once with complete media and resuspended in PBS for immunostaining. Cells were incubated with FITC anti-CD18 (clone 6.7, BD Biosciences, 1:100) for 30 minutes at 4C in the dark then washed once with PBS. Fluorescence minus one controls were prepared for CD18 conditions. Data were acquired on a BD Fortessa cytometer and exported to FlowJo (BD Biosciences) for analysis and graphed with Prism 8.0 (GraphPad).

### Lysate preparation and Western blots

Cell lysates from 3 × 10^6^ cells were generated using RIPA Lysis and Extraction Buffer (Thermo Fisher Scientific; 89901) supplemented with 1X Halt protease inhibitor cocktail (Thermo Fisher Scientific; 78443). Samples were incubated at 95°C with NuPAGE sample reducing agent (Thermo Fisher Scientific; NP0004) and NuPAGE LDS sample buffer (Thermo Fisher Scientific; NP0007) for 10 minutes. 5 × 10^5^ cell equivalents per well were loaded into a NuPAGE 4-12% Bis-Tris density gradient gel (Thermo Fisher Scientific; NP0432) and ran at a constant 150V for 80 minutes. Separated proteins were transferred onto nitrocellulose membranes at a constant 0.2A for 90 minutes. The nitrocellulose membranes were then blocked with 5% nonfat milk in PBS 0.05% Tween-20 for 60 minutes at 4°C. Nitrocellulose membranes were incubated overnight at 4°C with primary antibodies in 5% (w/v) BSA in PBS 0.05% Tween 20 at the following dilutions: 1:1000 anti-CD56 (clone 123C3; Cell Signaling Technology; 3576) and 1:4000 anti-actin (polyclonal; Sigma-Aldrich; A2066) as a loading control. Membranes were washed with 0.5M NaCl in PBS 0.05% Tween 20. Primary antibodies were probed with IRDye 680RD Goat anti-mouse IgG (Licor Biosciences; 926-68070) or IRDye 800CW goat anti-rabbit IgG (Li-COR Biosciences; 926-32211) secondary antibodies 1:10,000 for 1 hour at room temperature. Nitrocellulose membranes were imaged using the Odyssey CLx imaging system (Li-COR Biosciences).

### Sample preparation for live cell imaging

Glass chamber slides were coated with 0.001% Poly L-Lysine (PLL; Sigma P4707) for 1 hour at room temperature, washed 5 times with PBS, and then coated with a solution of PBS containing 5 µg/ml of fc-ICAM-1 (RD; 720-IC-200), anti-CD18 (purified from hybridoma; clone TS1/18), or 5 µg/ml Control:Fc (Enzo Life Science; ALX-203-004-C050) overnight at 4C. Each well was then washed 3 times with cell culture media. WT or CD56-KO NK92 cells were counted, centrifuged at 300 x*g* for 3 minutes, washed with media, and centrifuged for a further 3 minutes at 300 x*g* before being resuspended in media. 5 × 10^4^ cells were added to each well and cells were incubated at 37°C in the presence of 5% CO2 on the microscope for 1 hour to adhere cells to the surface prior to imaging.

### Confocal microscopy

WT and CD56-KO NK92 cells were incubated for 30 min (antibody coating) or 45 min (Fc-ICAM coating) at 37°C and 5% CO2 on #1.5 glass coverslips that had been pre-coated with 10 µg/ml anti-CD18 (clone IB4) and anti-NKp30 (clone P30-15, Biolegend; 325202) or 5 µg/ml Fc-ICAM-1 (Enzo Life Science; ALX-203-004-C050). NK cells were fixed and permeabilized with CytoFix/CytoPerm (BD Biosciences; 554714) at room temperature for 20 mins. Fixative was removed and coverslips were rinsed three times with 150 µl PBS 1% BSA 0.1% saponin. The following antibodies and reagents were used for fixed cell staining at 1:100 dilutions: phalloidin Alexa Fluor 568 (Thermo Fisher Scientific A12380), CD56 Alexa Fluor 647 (clone HCD56; Biolegend; 318314), CD44 Alexa Fluor 488 (clone C44Mab-5; Biolegend; 397508), open conformation LFA-1 Alexa Fluor 488 (clone m24; Biolegend; 363404), phospho-Pyk2 (Tyr402) (polyclonal; Abcam; ab4800), phospho-Pyk2 (Tyr881) (polyclonal; Abcam; ab4801), phospho-ezrin/radixin/moesin (Thr558) (polyclonal; Thermo Fisher Scientific; PA5-38679), phospho-Fyn (Tyr530) (polyclonal; Thermo Fisher Scientific; PA5-104756), goat anti-mouse IgG (H+L) cross-adsorbed secondary antibody Alexa Fluor 488 (polyclonal; Thermo Fisher Scientific; A-11001), goat anti-rabbit IgG (H+L) cross-adsorbed secondary antibody Alexa Fluor 488 (polyclonal; Thermo Fisher Scientific; A-11008). Coverslips were mounted on slides with ProLong Glass antifade reagent (ThermoFisher Scientific; P36934). Images were acquired with a 100X 1.46 NA objective on a Zeiss AxioObserver Z1 microscope stand equipped with a Yokogawa W1 spinning disk. Illumination was by solid state laser and detection by Prime 95B sCMOS camera. Data were acquired in SlideBook software (Version 6, Intelligent Imaging Innovations) and exported as TIFF files for further analysis. Live actin imaging was performed using confocal imaging on the same system. Timelapse images of a single axial plane close to the coverslip were acquired every 1 second for 120 seconds per cell. Laser power was set at 20% with integration time of 50 ms using a 561 nm laser and associated filters. Low resolution phase contrast imaging was performed using the same microscope with a 20X 0.5 NA magnification lens. Images were captured every 10 seconds for 30 minutes. For cytochalasin D treatment, cells were incubated on functionalized surfaces for 20 min prior to addition of cytochalasin D to a final concentration of 2 µM then incubated for 10 min before imaging. An environmental chamber maintained consistent 5% CO2 and 37°C (OKOlab).

### Structured illumination microscopy

WT and CD56-KO NK92 cells were prepared as described for confocal microscopy. Images were acquired on a GE Deltavision OMX SR equipped with a 60X 1.42 NA APO objective and 3 PCO Chip sCMOS cameras. 3D images were captured with 0.125 µm steps with a pixel size of 0.079 µm. SIM images were reconstructed with GE SoftWoRX software using 3 orientations and 5 phase shifts with a Wiener filter constant of 0.005 and negative values not discarded.

### Lattice lightsheet microscopy

WT and CD56-KO Life-Act mScarlet NK92 cells were resuspended in low serum NK92 media and then mixed with collagen that had been neutralized for a final collagen concentration of 1.6 mg/ml. Cells and collagen were incubated for 90 minutes in an Ibidi microchannel slide with #1.5 glass bottom. Cells were imaged on a Zeiss Lattice Lightsheet microscope with environmental control maintaining 37°C and 5% CO2. Cells were imaged at 20s intervals for 20-30 minutes. Deconvolution by constrained iterative algorithm and deskewing was performed using Zen (Zeiss Microsystems).

### Cell tracking analysis

Custom Fiji macros were used to organize TIFF stacks exported from SlideBook software into time-discriminated TIFF image sequences in replicate and condition folder hierarchy (21). Images were segmented using the Cyto2 trained network provided by Cellpose (19) using a classification object diameter of 30 pixels for data acquired at 20X magnification and 150 pixels for data acquired at 100X magnification and was run from the Anaconda command line. Bayesian Tracker (20) was used to track segmented cells between frames, and was run through a custom, generalizable Jupyter Notebook script (21). A HDF5 file containing segmented masks and tracks for each cell was output for each TIFF stack replicate and saved in the folder hierarchy. Custom Python functions were used to make 30 separate shape, migration, and clustering measurements per timepoint per cell and save these in a large Pandas dataframe containing all conditions and all replicates for each experiment (21). Migration measurements were mostly made between adjacent frames, but some measurements were normalized over 60 second time windows to compare tracks of different lengths; namely Euclidean distance, cumulative distance, and arrest coefficient. The radius r used for Ripley’s L(r) was 13.5 µm, 1.5X the average diameter of a single cell. Data was filtered to exclude objects smaller than 10 µm^2^ and cells tracked for fewer than 12 frames (120 seconds) for 20X data and 120 frames (120 seconds) for 100X data as they were assumed to be floating debris.

### Microscopy data processing and analysis

Line profiles were generated on cells that had polarized in response to incubation on ICAM-1. Polarized cells were identified by the accumulation of CD44, and line profiles were generated while blinded to CD56 distribution to avoid selection bias of cells for analysis. Line profiles were generated in both channels from uropod straight to the front of the cell in Fiji then normalized to the maximum value for each condition. Line profiles from 20 cells were plotted.

Analysis of m24 foci was performed in Fiji (36). Single slices at the plane of the glass were selected using the actin (phalloidin) channel. Thresholding was performed based on a value that excluded background from outside the cell and the threshold value was consistent between all cells in all conditions from a given experiment. Measure Particles was used with a size exclusion lower limit of 0.01 µm^2^ and no upper limit or circularity restriction. Area of individual foci and frequencies (numbers of foci per cell) were plotted in Prism (GraphPad Software).

### Plotting and statistical analysis for live cell imaging

Due to a high volume of data, p-values from automated live cell tracking were in some cases very small. We sought to display the effect size to better understand comparisons between conditions. Plots of Difference (37) were adapted for Python and Matlab (from R) and are used here to calculate the effect size distribution. Briefly, each condition is resampled 1000X to calculate a new distribution of mean values by bootstrapping. Each mean value is subtracted from the mean value of a user supplied control condition, resulting in a new distribution of differences representing the ‘effect size’ which is plotted as a distribution next to the data. The mean effect size and confidence intervals and their distance/overlap with the ‘control’ condition indicate the biological and statistical significance of the change respectively. P-values were calculated using the randomization method, making no assumption about the underlying data distribution (38). Superplots of Data (39) were also adapted for Python and Matlab (from R) and used to transparently display data from individual replicates and individual conditions.

### Optical flow analysis

Images were exported from Slidebook software as TIFF stacks. Fiji was used to pre-select cells with average intensity > 300, indicating the presence of enough tagged actin on which to perform optical flow analysis. Fiji was also used to order the files into folders for batch analysis. To measure the magnitude and directionality of actin flow, we adapted an algorithm based on the Lucas-Kanade method (40), which calculates vectors for flows of ‘like’ pixel intensities detected between subsequent frames. The x/y vector components are used to generate vector maps encoding the magnitude and directionality of flow. Masks based on the cell contour and a reliability matrix for each vector based on properties of the underlying image was used to remove noise. A lower reliability threshold of 0.01 times the maximum reliability value for each individual cell was used to select reliable vectors. The number of pixels with no reliable vectors or vectors below a lower magnitude threshold of 0.5 µm/min were classed as immobile and were divided by the total number of pixels within the cell to give the immobile fraction per time point. The distribution of flow vectors for overlapping time windows of 10 frames (10 seconds) were fit to a von Mises distribution, and the kappa parameter was used to indicate the persistence of motion of flow in a single direction (41). Higher kappa values indicated more persistently directional flow. Batch analysis of cells in folders and output of all plots is possible by changing a single directory line, pixel size, and time between frames in the code provided by Dr. Arpita Upadhyaya that is available at https://zenodo.org/record/8221517.

## Code availability

Optical flow analysis software was kindly provided by Dr. Ivan Rey-Suarez (Upadhyaya Lab) and was adapted here for batch analysis and plotting of actin flow speed, Von Mises concentration of motion, and calculation of the immobile fraction. This version of the software is available at https://zenodo.org/record/8221517. Calculations of migration characteristics, cell clustering and morphology per cell per time point, including Plots of Difference (42), TimePlots of Difference and SuperPlots (39, 43) were created using cellPLATO (21) with code available at https://zenodo.org/records/8096717.

## ACKNOWLEDGEMENTS

We thank Drs. Arpita Upadhyaya and Ivan Rey-Suarez for sharing code for PIV analysis and Alfred Kibowen and Drs. Chris Bjornsson (Zeiss Microsystems) and Abhishek Kumar (MBL) for technical assistance with lattice lightsheet microscopy at the Marine Biological Laboratory. Part of this work was initiated during the Lightsheet Fluorescence Microscopy (LSFM) course and workshop at the MBL and supported by a Whitman Fellowship to EMM. Research reported in this publication was supported in part by the National Institute of General Medical Sciences of the National Institutes of Health under award number R01GM148504 to EMM.

