## Supplemental Figures for "CD56/NCAM mediates cell migration of human NK cells by promoting integrin-mediated adhesion turnover"

Supplementary Figure 1

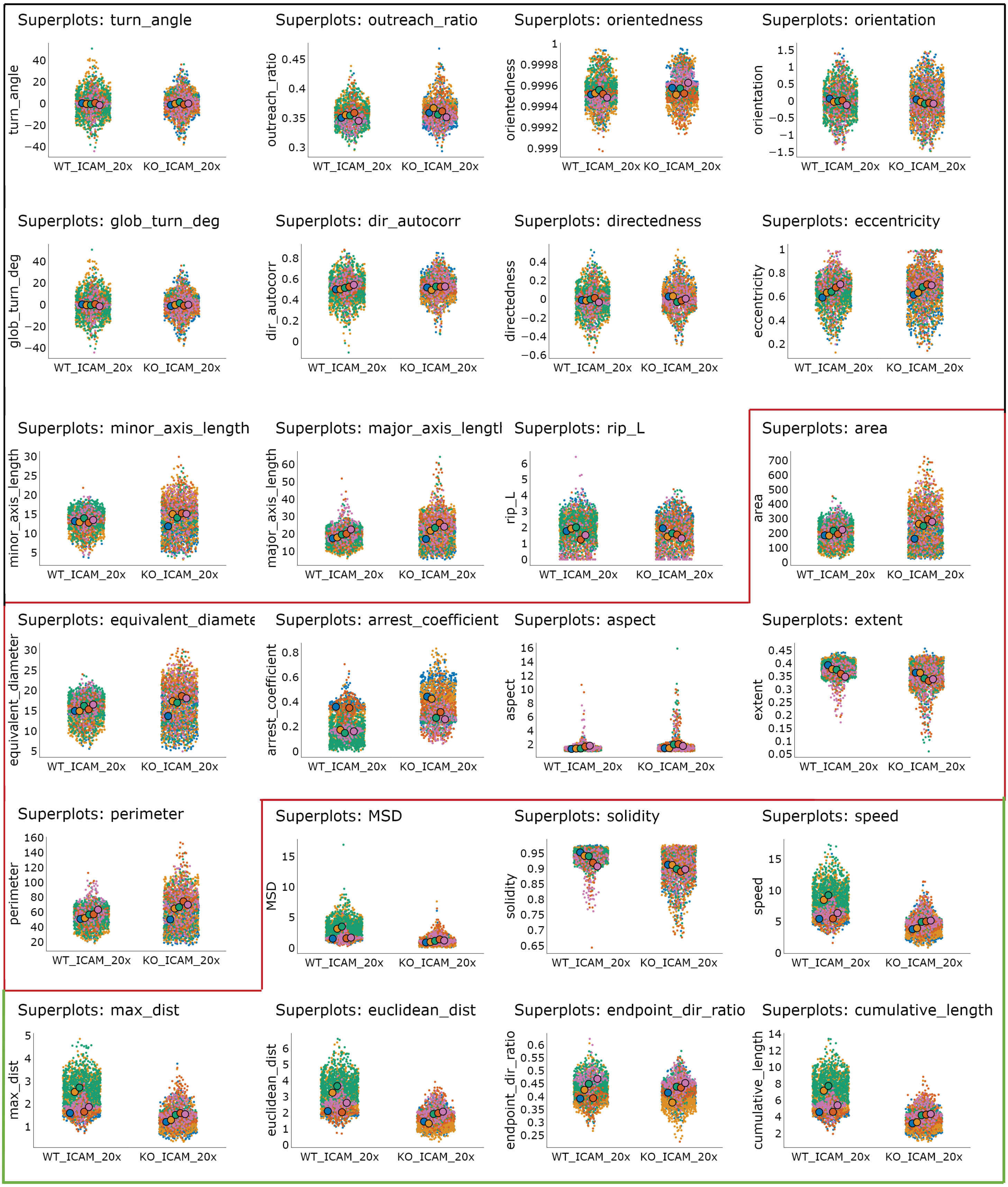


**Supplementary Figure 1. All measurements taken for WT and CD56-KO NK92 migrating on glass coated with ICAM-1.** SuperPlots show all the data (small points) and mean values (large points) from separate replicates denoted by colour. Metrics outlined in black do not change between CD56 proficient and deficient cells; red outline: metrics increased in CD56 deficient cells; green outline: metrics decreased in CD56 deficient cells. Units are all μm, μm2 and μm/min. n=5 experimental replicates per condition, n= 3946 cells.

Supplementary Figure 2

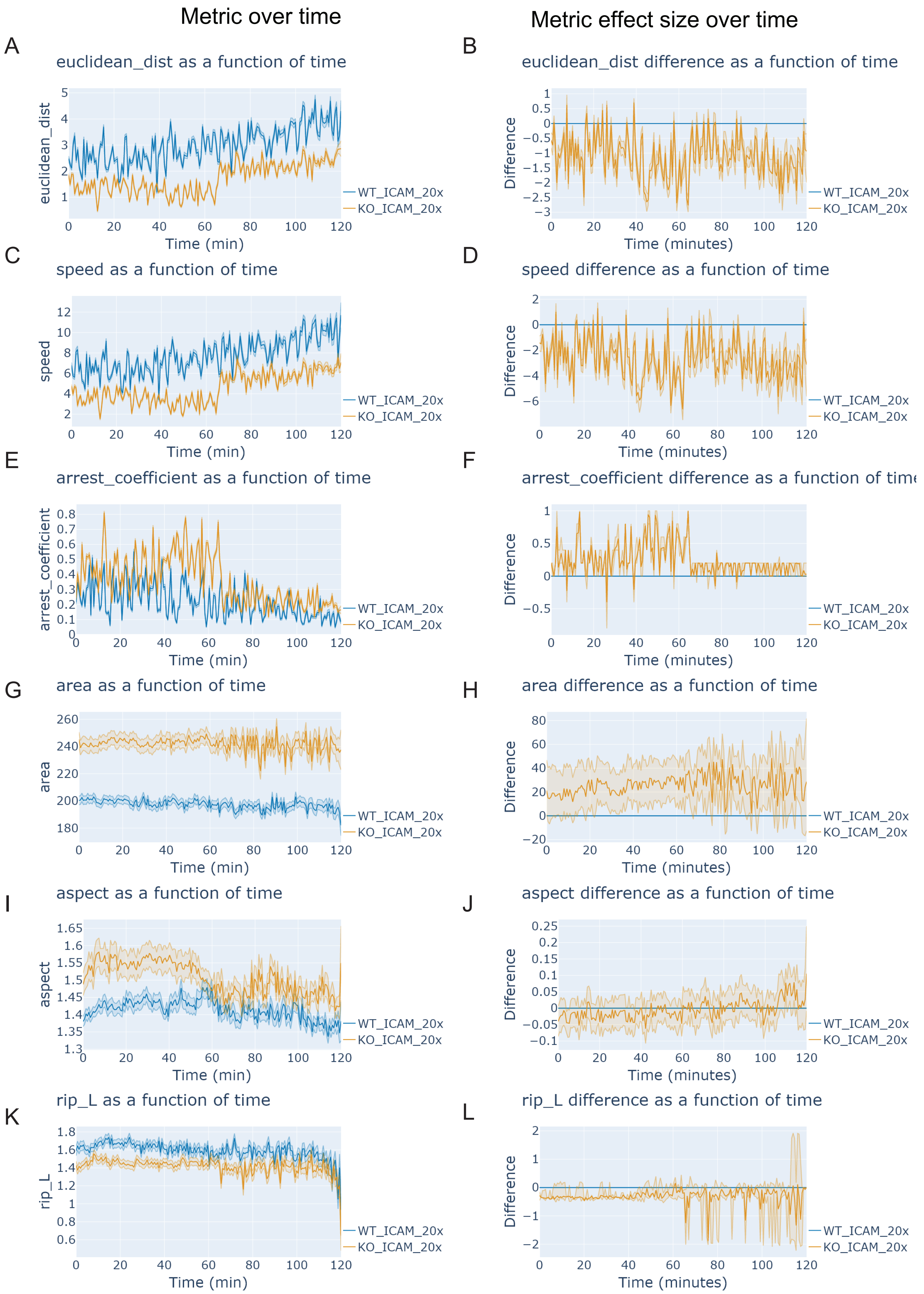


**Supplementary Figure 2: Time plots of difference show that migration, morphology, and clustering metrics remain constant over time**. A) Mean Euclidean distance (thick lines) and confidence intervals (transparent regions) over time, B) effect size over time for Euclidean distance; blue line at 0 = control condition NK92 cells, orange line = NK92 CD56-KO cells. C and D) Cell Speed and effect size, E and F) Arrest coefficient and effect size; G and H) Area and effect size; I and J) Aspect ratio and effect size, K and L) Ripley's L (clusteredness) and effect size.

Supplementary Figure 3

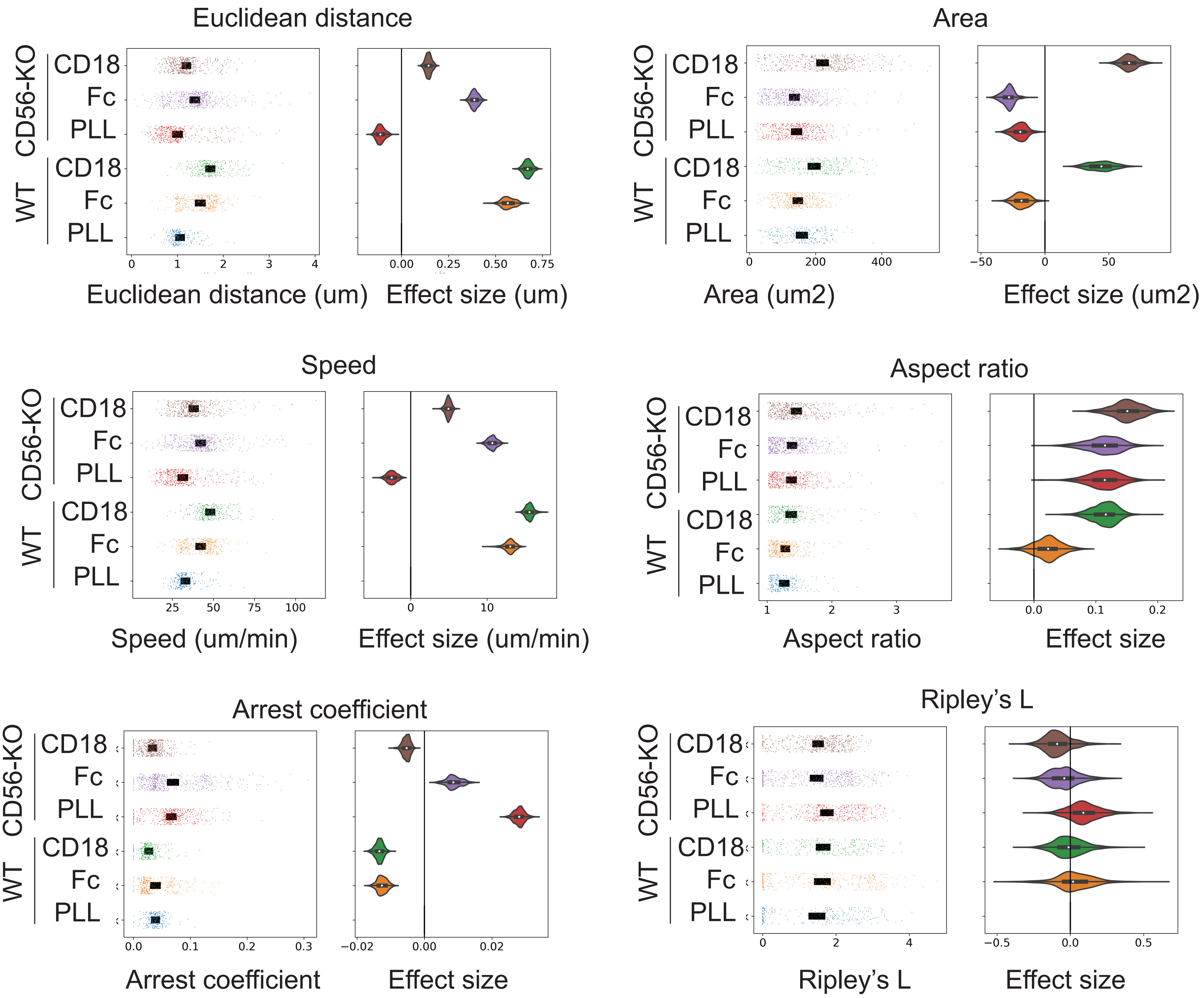


**Supplementary Figure 3: NK92 WT and CD56-KO cell behavior changes on Fc-, PLL and anti-CD18 coated glass surfaces.** WT or CD56-KO NK92 cells were incubated on glass slides coated with a human Fc, PLL or anti-CD18 and imaged by phase contrast for 30 minutes with 10 seconds between frames. Plots of difference show the distribution of the data (left pane) and effect size (right pane) for metrics related to migration, morphology, and clustering: A) Euclidean distance, B) Area, C) Speed, D) Aspect ratio, E) Arrest Coefficient and F) Ripley's L (clusteredness). n=2 experimental replicates per condition, n= 2233 cells. Cells included in the analysis were filtered to include at least 2 minutes of tracking and a minimal area of 10 μm$^2$ to exclude debris.

Supplementary Figure 4

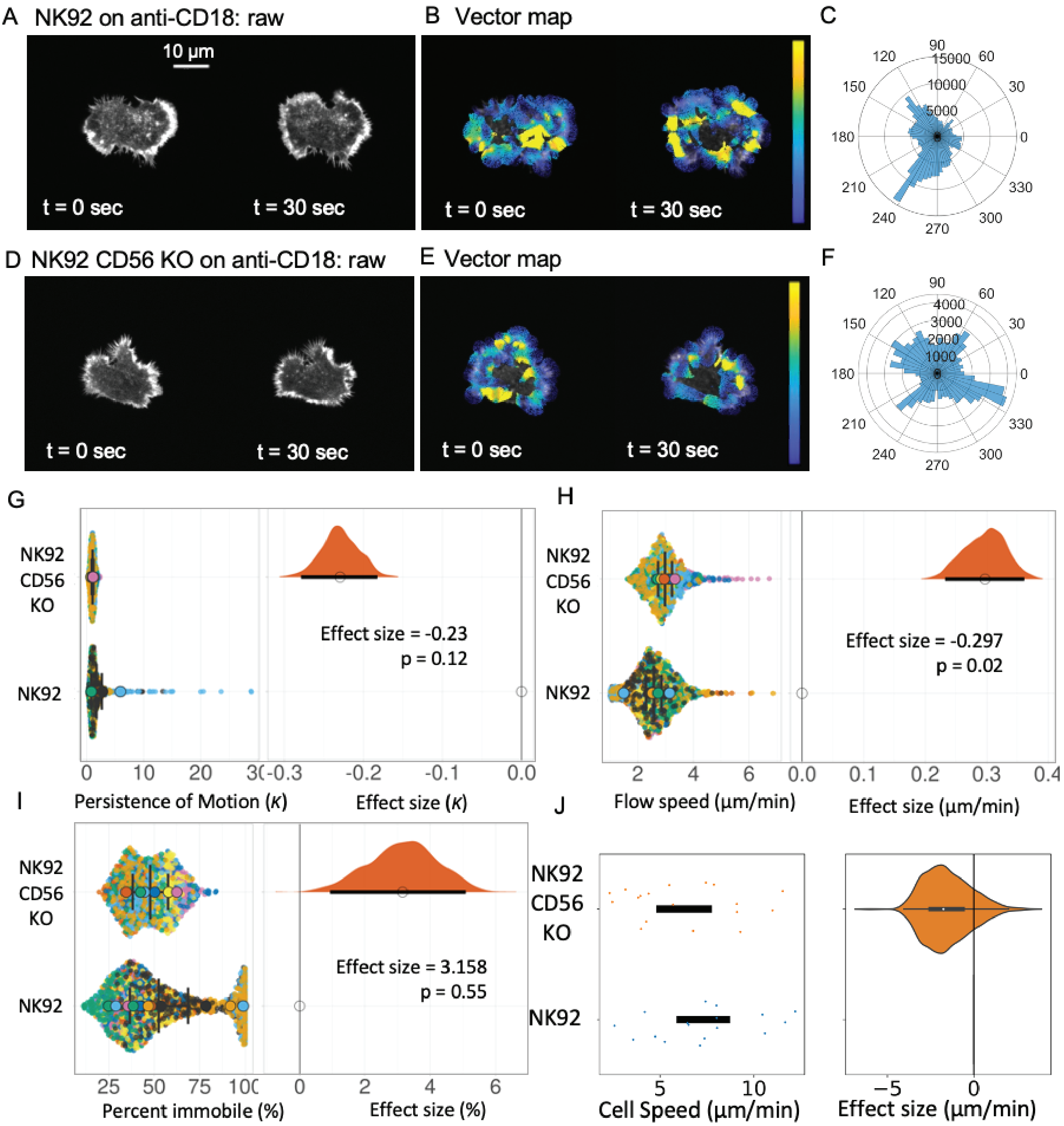


**Supplementary Figure 4. Actin flow speed and immobile fraction are unaffected in NK92 CD56-KO cells spreading in response to CD18 antibody binding.** WT or CD56-KO NK92 cells expressing LifeAct mScarlet were incubated on glass slides coated with anti-CD18 and imaged at 100X magnification for 2 minutes per field of view, with 1 second between frames using a 561 nm laser. A) Actin in NK92 WT cells, example frames from a timelapse movie are shown (0 and 30 seconds). B) Lucas-Kanade PIV vector field (warm colors = high magnitude, cold = low), and C) polar histogram of directionality. D, E and F) raw images, vector field and polar histogram of directionality for NK92 CD56 KO cells. Plots of difference showing G) the concentration of motion of flow (κ from the von Mises fit) across 10 second overlapping time windows illustrating change in flow direction, H) actin flow speed, I) percent of actin that was immobile, J) cell speed of migration. n=12 and 7 experimental replicates per condition respectively, n= 19 cells analyzed.

Supplementary Figure 5

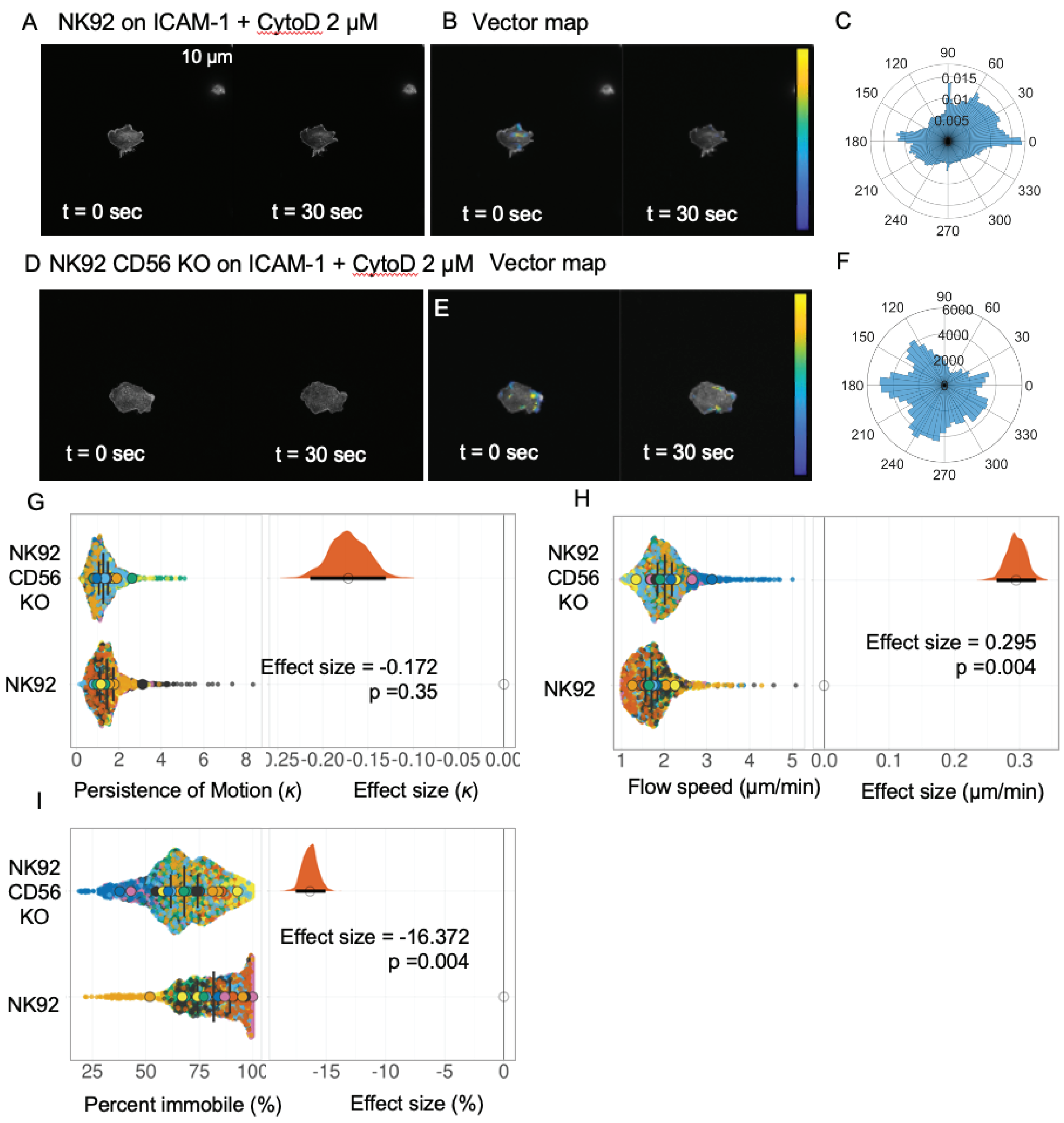


**Supplementary Figure 5. Actin flow speed, immobile fraction and Von Mises concentration of motion in Cytochalasin D treated cells on ICAM-1.** WT or CD56-KO NK92 cells expressing LifeAct mScarlet were incubated on glass slides coated with Fc-ICAM-1 and were imaged at 100X magnification for 2 minutes per field of view with 1 second between frames using a 561 nm laser. A) Actin in NK92 WT cells, example frames from a timelapse movie are shown (0 and 30 seconds). B) Lucas-Kanade PIV vector field (warm colors = high magnitude, cold = low), and C) polar histogram of directionality. D, E and F) raw images, vector field and polar histogram of directionality for NK92 CD56 KO cells. Plots of difference showing G) the concentration of motion of flow (κ from the von Mises fit) across 10 second overlapping time windows illustrating change in flow direction, H) actin flow speed, I) percent of actin that was immobile. n=15 and 21 experimental replicates per condition respectively, n= 36 cells analyzed.

Supplementary Figure 6

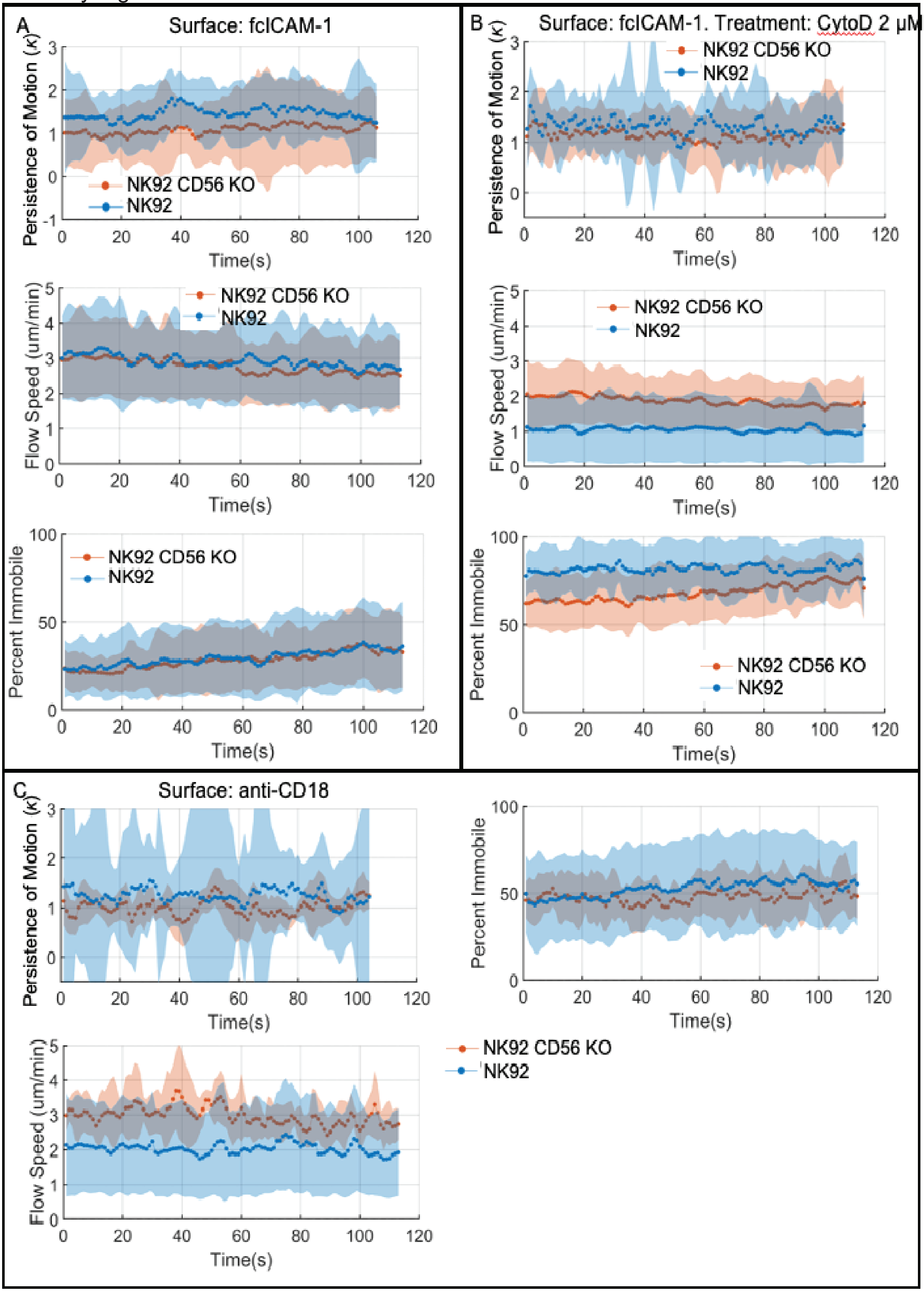


**Supplementary Figure 6. TimePlots of Difference of NK92 cells migrating on ICAM-1 or anti-CD18 surfaces with and without Cytochalasin D.** WT or CD56-KO NK92 cells expressing LifeAct mScarlet were incubated on glass slides coated with fc-ICAM-1 and were imaged at 100X magnification for 2 minutes per field of view with 1 second between frames using a 561 nm laser. A) NK92 cell timelapse frames (top) and PIV analysis (bottom), and B) the same for NK92 CD56-KO cells. Hot colors indicate fast flow and cold colors indicate slow flow. C) Actin flow was calculated using the Lucas-Kanade PIV method. D) Percentage of the cellular actin not flowing was calculated from the cell masks used for PIV. E) The concentration of motion of flow (κ from the von mises fit) across 10 second overlapping time windows illustrates change in flow direction. F) Cell speed, effect size and area plotted for cells imaged at 100X in (G). n=30 and 25 experimental replicates per condition respectively, n= 92 cells.

**Supplementary movies (available at https://zenodo.org/records/10180692)**

**Movie 1.** Representative fields of view from experiments shown in Fig. 2. WT NK92 cells were incubated on ICAM-1 coated glass and imaged by live cell phase contrast microscopy at 10 second intervals for 30 minutes. Cell masks generated by segmentation are shown on the right.

**Movie 2.** Representative fields of view from experiments shown in Fig. 2. CD56-KO NK92 cells were incubated on ICAM-1 coated glass and imaged by live cell phase contrast microscopy at 10 second intervals for 30 minutes. Cell masks generated by segmentation are shown on the right.

**Movie 3.** Representative cells from experiments shown in Fig. 3. WT NK92 cells expressing LifeAct mScarlet were incubated on ICAM-1 coated glass and imaged by live cell confocal microscopy at 100X magnification for 2 minutes per field of view with 1 second between frames using a 561 nm laser.

**Movie 4.** Representative cells from experiments shown in Fig. 3. CD56-KO NK92 cells expressing LifeAct mScarlet were incubated on ICAM-1 coated glass and imaged by live cell confocal microscopy at 100X magnification for 2 minutes per field of view with 1 second between frames using a 561 nm laser.

**Movie 5.** Representative cells from experiments shown in Supp. Fig. 4. WT NK92 cells expressing LifeAct mScarlet were incubated on anti-CD18 coated glass and imaged by live cell confocal microscopy at 100X magnification for 2 minutes per field of view with 1 second between frames using a 561 nm laser.

**Movie 6.** Representative cells from experiments shown in Supp. Fig. 4. CD56-KO NK92 cells expressing LifeAct mScarlet were incubated on anti-CD18 coated glass and imaged by live cell confocal microscopy at 100X magnification for 2 minutes per field of view with 1 second between frames using a 561 nm laser.

**Movie 7.** Lattice lightsheet imaging of WT NK92 cells expressing LifeAct mScarlet embedded in 1.6 mg/ml collagen and imaged live every 40 sec.

**Movie 8.** Lattice lightsheet imaging of CD56-KO NK92 cells expressing LifeAct mScarlet embedded in 1.6 mg/ml collagen and imaged live every 40 sec.
